# Evaluating the Effects of Environmental Disturbances and Pesticide Mixtures on N-cycle related Soil Microbial Endpoints

**DOI:** 10.1101/2024.01.22.576671

**Authors:** Camilla Drocco, Anja Coors, Marion Devers-Lamrani, Fabrice Martin-Laurent, Nadine Rouard, Aymé Spor

**Affiliations:** Agroécologie, INRAE, Institut Agro, Université de Bourgogne, Dijon, France; ECT Oekotoxikologie GmbH, Flörsheim/Main, Germany

**Keywords:** ecotoxicology, global change disturbance, heat stress, Nitrogen cycle, pesticide mixture, bacterial communities

## Abstract

Pesticides are widely used in conventional agriculture, either applied separately or in combination during the culture cycle. Due to their occurrence and persistence in soils, pesticide residues may have an impact on soil microbial communities and on supported ecosystem services. In this regard, the EFSA (European Food Safety Authority) recently published a scientific opinion inciting to change pesticide risk assessment to better protect soil microbe-mediated processes. Climate change is another major concern for all living organisms including soil microbial community stability. Extreme climatic events, such as heat waves or heavy rainfalls, are becoming more and more frequent and their impact on soil microbial diversity and functions have already been demonstrated.

The objectives of this study were to evaluate the effects of temperature and humidity disturbances and pesticide active ingredients exposure on soil microbial community structure and functions. To this end, 250 soil microcosms were exposed to either a heat disturbance, a high humidity to mimic heavy rain, or no environmental disturbance. After three days of recovery, soil microcosms were treated with different active ingredients: clopyralid (herbicide), cypermethrin (insecticide) and pyraclostrobin (fungicide). The treatments were applied alone or in combination at 1x or 10x of the agronomical dose. We then evaluated the effects of the disturbances and the active ingredients on various microbial endpoints related to the diversity and the structure of soil microbial communities, and with a specific focus on microbial guilds involved in nitrification.

Overall, we demonstrated that the impact of environmental disturbances applied to soil microcosms, especially heat, on microbial endpoints was stronger than that of the active ingredients applied alone or in combinations. Compounded effects of environmental disturbances and active ingredients were detected, but sparsely and were of small scale for the chosen pesticides and applied doses.

## INTRODUCTION

Soil is a non-renewable resource at human life span that carries out several ecosystem services supporting the life of many organisms: it sustains food and fibers production, stores CO_2_ and water, and helps in disease suppression (Dominati et al., 2010; Aislabie and Deslippe, 2013; Moebius-Clune et al., 2016). Soil microbial communities are major drivers of nutrient cycles that sustain plant growth and productivity. The nitrogen cycle (N-cycle), which regulates the availability of N in soil, is carried out by specific microbial guilds (Dominati et al., 2010; Aislabie and Deslippe, 2013; Whitman et al., 1998; Singh, 2015). Indeed, the N_2_ in the atmosphere is not available for direct use by plants, while symbiotic nitrogen-fixing bacteria can fix it into plants as ammonia that is then used to build amino acids and proteins (Moebius-Clune et al., 2016). The ammonia oxidizing bacteria and archaea take part to the nitrification converting the NH_4_^+^ derived from organic nitrogen into NO_2_^−^ (Lehtovirta-Morley, 2018), while the newly discovered comammox can transform ammonia directly into NO_3_ ^−^ (Koch et al., 2019). The inverse process from NO_2_^−^ to N_2_ is performed by the denitrifiers or anammox bacteria (Kuypers et al., 2018). Overall, the N-cycle involves a wide diversity of microbial taxa that are important to ensure its functioning all along seasonal changes of soil temperature and humidity.

Climate change and extreme climatic events can induce shifts in microbial community composition and taxa abundance (Barnard et al., 2013; Jurburg et al., 2017a; Mooshammer et al., 2017; Calderón et al., 2018; Meisner et al., 2018), with effects on nutrient cycles (Philippot et al., 2013) and ecosystem resilience (Jurburg et al., 2017a). Frey et al. (2008) in their long-term experiment in mixed forest concluded that 12 years of soil heating reduced total microbial biomass, and induced a shift in community composition, with a negative impact on substrate use. The N-cycle is also deeply influenced by temperature (Calderón et al., 2018; Mooshammer et al., 2017; Sahrawat, 2008; Szukics et al., 2010), and the perturbation of one of the steps, *e. g.* nitrification, poses challenges for nitrogen supply in the soil, leading to the disruption of nitrogen availability.

Pesticides are commonly applied in conventional agriculture and in integrated pest management to ensure an adequate production of food and fibres. According to the Food and Agriculture Organization (FAO, 2023), 354’082 tonnes of pesticides were sprayed for agricultural use in the EU in 2021, of which only a part of the applied compounds gets to the target. The rest (30-50%) can reach the soil environment (Rodríguez Eugenio et al., 2018) where it may impact non-target organisms and transfer to other environmental compartments, including water resources. Several studies have already documented that pesticide residues can impact soil microbial community leading to *i*) the emergence of pesticide-degrading strains able to use pesticide as carbon source for their growth, or *ii*) the alteration of microbial abundance and diversity and possibly affecting ecosystems stability and their resilience (Karas et al., 2018; Kumar et al., 2019; Puglisi, 2012; Storck et al., 2018; Zabaloy et al., 2010). During the entire crop cycle, often more than one pesticide is applied at the same field. Persistent residues of active ingredients as well as their transformation products accumulate to a cocktail of chemicals to which soil living organisms are exposed (Geissen et al., 2021; Tang and Maggi, 2021). The overall ecotoxicological impact resulting from the exposure of non-target organisms to chemical cocktails is the result of the combination of additive, antagonistic or synergistic effects, that until now have been rarely studied with regard to soil microorganisms. Cedergreen (2014) reviewed the available literature on chemical mixtures, concluding that additive interactions generally dominate whereas evidences for synergistic interactions were limited to cholinesterase inhibitors (insecticide) and cytochrome P450 inhibitors (fungicide). Toxicity endpoints of aquatic organisms dominated this review (Cedergreen 2014), while microbial endpoints were underrepresented. Hence, there is a strong need for a better understanding of the impact of accumulated pesticide residues particularly on the soil microbial community.

Ecosystems are usually exposed to more than one disturbance, that act independently or influence each other, with different effects on the ecological recovery of the ecosystem functions. Very often, a legacy effect of disturbances (Paine et al., 1998) can be observed and might decrease the ability of species to resist and then recover to subsequent perturbations. Jurburg et al. (2017b) demonstrated that a microbial community needed at least 29 days to recover from a heat shock. In addition, the disturbance history (heat, and heat or cold shock) affects the microbial community response to the subsequent perturbations (Jurburg et al., 2017a). Extensive works have been done to assess the effect of factors mimicking extreme climate events (Bardgett and Caruso, 2020; Calderón et al., 2018; Meisner et al., 2018; Mooshammer et al., 2017; Philippot et al., 2021), but up to now, only a few studies have considered the potential combined effect of environmental disturbances and pesticides on soil microbial communities (Franco-Andreu et al., 2016; Pesaro et al., 2004).

Within this context, the objectives of the present work are, therefore, to compare effects of global change-related environmental disturbances and impacts of pesticidal active ingredients on soil microbial community structure and functioning and to evaluate their compounded effects. We hypothesize that, as compared to undisturbed soil *i*) extreme climatic events, such as elevated temperature and heavy rainfall will change microbial community’s structure and functioning; *ii*) subsequent exposure to different pesticidal active ingredients applied alone or in combination may lead to compounded effects on the structure and functioning of some key microbial community members, namely guilds involved in N-cycle.

## MATERIAL AND METHODS

### Soil sampling and microcosms set up

Soil from the surface layer (0-20 cm depth) was sampled from Sayens field site located in the vicinity of the INRAE experimental farm in Bretenière, France (47° 14’ 03.1” N, 5° 06’ 37.9” E) during August 2021. This soil is silty clay loam, characterized by 34.6% clay, 59.0% silt, and 6.3% sand, with an organic carbon content of 27.3 g.kg^−1^ and a nitrogen content of 2,7 g.kg^−1^. The water holding capacity (WHC) was 64. The soil was homogenized and sieved to 4 mm and stored at 4°C until further use. The soil humidity after sieving was 26% which corresponds to ∼ 40% of the WHC. The fresh soil was divided into 250 microcosms, each containing 63 g fresh soil (corresponding to 50 g dry soil), and incubated at room temperature for 6 days prior to exposure, or not, to environmental disturbance.

### Environmental disturbances and active ingredients application

We studied two different environmental disturbance factors: elevated temperature (heat) and heavy rainfall (high humidity). Respectively, 80 soil microcosms were subjected to the heat disturbance, 80 to high humidity and 80 received no environmental disturbance (control). Heat and high humidity disturbances were applied over a window of 7 days. The heat disturbance consisted in two periods of 30 h at 42°C, separated by a period of 40 h at room temperature. At the end of the disturbance window, the humidity was adjusted to 22%. The high humidity disturbance consisted in watering soil microcosms to reach 35% of humidity, followed by an evaporation period of 7 days. Three days after the end of the disturbance window, *i. e.* at day 19 of the experiment, five microcosms per each disturbance condition tested (heat, high humidity and control) were treated with one of the three pesticides (technical material of active ingredients, a.i.) or with the mixture of the three a.i at 1x or 10x the agronomical dose. The pesticides were: 3,6-dichloropyridine-2-carboxylic acid (*i e.* clopyralid, pesticide analytical standard, CAS 1702-17-6, supplier: Sigma-Aldrich, purity ≥ 98.0%, agronomical dose: 125 g/ha corresponding to 4.5 µg/50 g dry soil), [cyano-(3-phenoxyphenyl)methyl] 3-(2,2-dichloroethenyl)-2,2-dimethylcyclopropane-1-carboxylate (*i. e.* cypermethrin, pesticide analytical standard, CAS 52315-07-8, supplier: Sigma-Aldrich, purity ≥ 90.0%, agronomical dose: 100 g/ha corresponding to 3.4 µg/50 g dry soil), and methyl *N*-[2-[[1-(4-chlorophenyl)pyrazol-3-yl]oxymethyl]phenyl]-*N*-methoxycarbamate (*i. e.* pyraclostrobin, pesticide analytical standard, CAS 175013-18-0, supplier: Sigma-Aldrich, purity ≥ 98.0%, agronomical dose: 200 g /ha corresponding to 7.1 µg/50 g dry soil). These three a.i are all constituents of commercial formulated products commonly used in agriculture, and can be all used in corn cropping during a growth cycle. Clopyralid was dissolved in water, and cypermethrin and pyraclostrobin were dissolved in acetone, and equal quantities of water (1 mL) and acetone (10 µL) a.i. solutions were applied to all microcosms with the respective treatment (single a.i or a.i. mixture). The solutions were applied onto the soil surface, without mixing the soil after application. In a similar manner, water and acetone were applied to 10 untreated control microcosms. Soil moisture was adjusted and kept to 26% in all microcosms. All the treated microcosms and the untreated controls were done in five independent replicates, for a total of 250 microcosms (Fig. 1).

**Figure 1.**
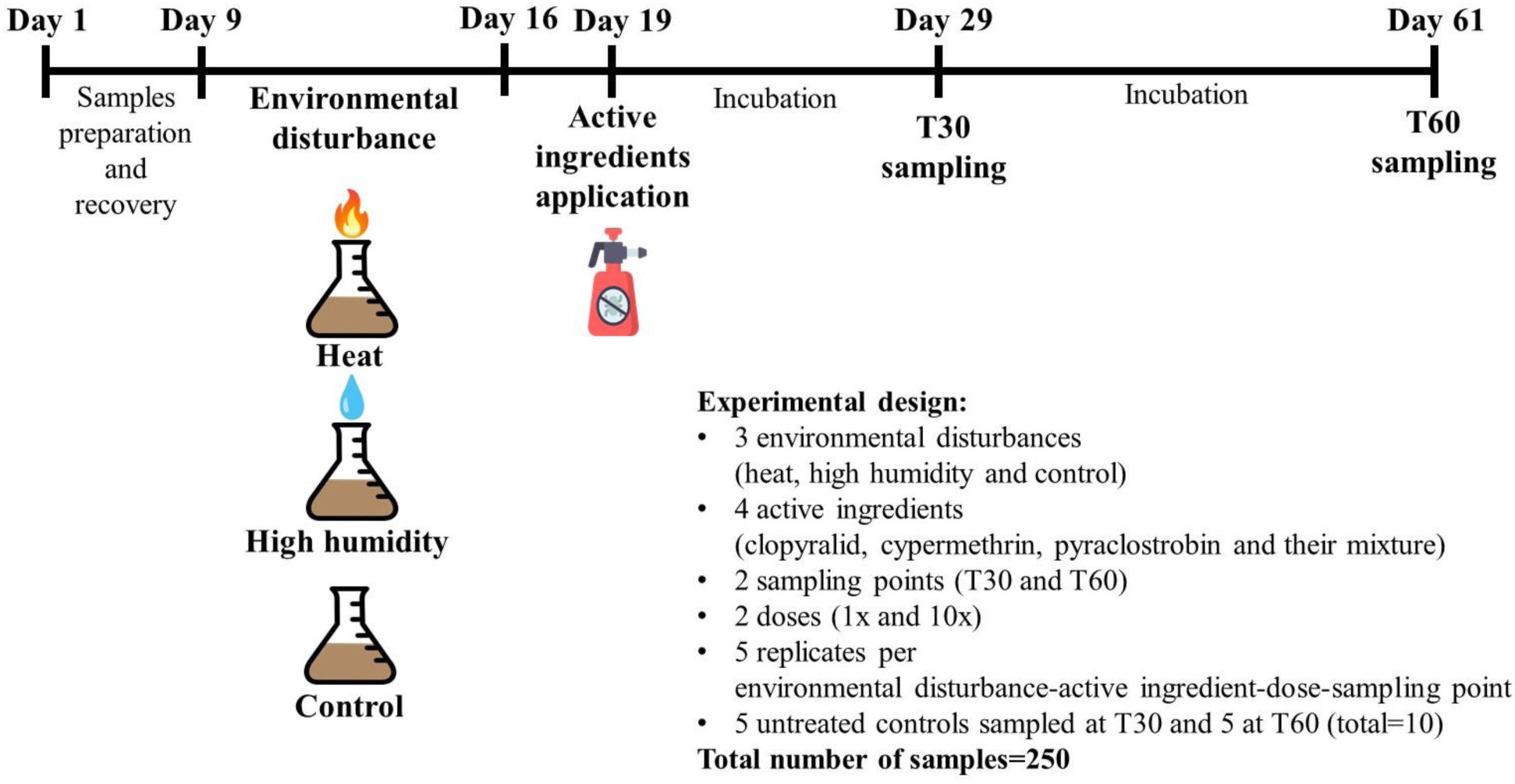
Graphic representation of the experimental timeline with a description of the experimental design.

From each treatment combination and from the untreated controls, five microcosms were destructively sampled at day 29 (T30: 10 days after a.i application), and the remaining five at day 61 (T60: 42 days after a.i application). The soil samples were processed fresh or stored at –20°C for further analyses.

### Nitrogen pools quantification

The nitrogen species NO_3_^−^ and NH_4_^+^ were extracted from 10 g (fresh weight) of the sampled soil using 50 ml of K_2_SO_4_ 0.5 M. The soil suspension was shaken for one hour, allowed to settle for two-three hours at room temperature, and then decanted. The supernatant was filtered and kept frozen until quantification. The quantification was performed by colorimetry according to ISO 14256-2:2005. Two blanks per series were included. The results of the NO_3_^−^ and NH_4_^+^ quantifications were expressed as mg of N/kg of dry soil.

### DNA extraction

DNA was extracted from 250 mg of dry weight soil using the DNeasy PowerSoil DNA extraction Kit (Qiagen, FR) following the kit’s instructions. The DNA was quantified with the Quant-iT™ dsDNA Assay Kit (Thermo Fischer Scientific, FR), and stored at –20°C until further use.

### Assessment of microbial community composition and diversity

Bacterial community composition and diversity were monitored via sequencing the 16S rDNA V3-V4 hypervariable region. We use a two-step PCR amplification method (Berry et al., 2011). In the first step, 25 amplification cycles were performed using the fusion primers U341F (5’-CCTACGGGRSGCAGCAG-3’) and 805R (5’-GACTACCAGGGTATCTAAT-3’) as described in Takahashi et al., (2014), with overhang adapters (forward: 5’-TCGTCGGCAGCGTCAGATGTGTATAAGAGACAG-3’, adapter: 5’-GTCTCGTG GGCTCGGAGATGTGTATAAGAGACAG-3’). The thermal cycling conditions of the first PCR were: 3 min at 94°C, 25*(30 sec at 94 °C, 30 sec at 55 °C, and 30 sec at 72 °C), and 10 min at 72°C. Duplicate of the first PCR were pooled and used in a second PCR, were the amplification linked a unique combination of multiplex primer pair to the overhang adapters for each sample. The thermal cycling conditions of the second PCR were: 3 min at 98°C, 8*(15 sec at 98°C, 30 sec at 55°C, 30 sec at 72°C), and 10 min at 72°C.

The amplicons from the second PCR were pooled and visualized in a 2% agarose gel for size and amplification check. They were then pooled and purified with the SequalPrep Normalization Plate kit (Invitrogen, Frederick, MD, USA). Sequencing was performed by GenoScreen (Lille, FR) on MiSeq (Illumina 2*250 bp) using the MiSeq reagent kit v2 (500 cycles).

### Quantification of microbial communities

The total abundances of bacteria (16S), fungi (ITS), and protists (18S), as well as the N-cycle microbial guilds were estimated through real-time quantitative PCR (qPCR). The 16S rDNA and ITS2 primers described by Muyzer et al. (1993) and White et al. (1990) were used to estimate the total 16S and ITS communities abundances, respectively. The 18S community abundance was estimated from the 18S rDNA using the primers EK-565F (5′-GCAGTTAAAAAGCTCGTAGT-3′) and 18S-EUK-1134-R–UnonMet (5′-TTTAAGTTTCAGCCTTGCG-3′) with cycling condition of 3 min at 95°C, 35*(10 sec at 95°C, 30 sec at 58 °C, 30 sec at 72 °C and 20 sec at 80°C). The ammonia oxidizing archaea (AOA) and bacteria (AOB) were quantified through gene *amoA* (Bru et al., 2011), comammox A (ComaA) and B (ComaB) were targeted with the clade A and B *amoA* genes (Pjevac et al., 2017), respectively.

The quantifications were carried out in a ViiA7^TM^ (Life Technologies, Carlsbad, CA, USA) with a reaction volume of 15 µL containing 7.5 µL of Takyon Master Mix (Eurogentec, Liège, BE), 1 µM of each primer, 250 ng of T4 gene 32 (MP Biomedicals, Santa Ana, CA, USA), 3 ng of DNA. Two independent replicates were used for each qPCR assay. Prior to qPCR analysis, an inhibition test was performed by mixing the soil DNA extracts with a control plasmid DNA (pGEM-T Easy Vector, Promega, Madison, WI, USA) or water. No inhibition was detected. The results from the quantification analysis were expressed as number of gene copies/ng of DNA.

### Bioinformatics analysis

Sequencing data were analyzed using an in-house developed Jupyter Notebooks (Kluyver et al., 2016) piping together different bioinformatics tools. Briefly, for 16S, R1 and R2 sequences were assembled using PEAR (Zhang et al., 2014) with default settings. Further quality checks were conducted using the QIIME pipeline (Caporaso et al., 2010) and short sequences were removed (< 400 bp for 16S). Reference based and de novo chimera detection, as well as clustering in OTUs were performed using VSEARCH (Rognes et al., 2016) and the SILVA v132 reference database. The identity thresholds were set at 94% for 16S rDNA data based on replicate sequencings of a bacterial mock community containing 40 bacterial species. Representative sequences for each OTU were aligned using infernal (Nawrocki and Eddy, 2013) and a 16S phylogenetic tree was constructed using FastTree (Price et al., 2009). Taxonomy was assigned using BLAST (Altschul et al., 1990) and the SILVA reference database v132 (Quast et al., 2013).

Diversity metrics, that is, Faith’s Phylogenetic Diversity (Faith, 1992), richness (observed species) and evenness (Simpson’s reciprocal index), describing the structure of microbial communities were calculated based on rarefied OTU tables (8300 sequences per sample). Weighted UniFrac distance matrices (Lozupone and Knight, 2005) were also computed to detect global variations in the composition of microbial communities.

### Statistical analysis of the microbial endpoints and of the nitrogen pools

The statistical analysis was carried out using RStudio statistical software version 4.2.1. The data were checked for homogeneity of variance with Levene’s test (*p* < 0.05). Data were log-transformed or cos-transformed when necessary and outliers were removed when detected (a maximum of 2 values were removed for any given variable).

The values from 16S, ITS, 18S, AOA, AOB, ComaA, ComaB abundances, NO_3_^−^ and NH_4_^+^ concentration, and α-diversity indices which are observed species (OS), Faith’s Phylogenetic Diversity (PD) and Simpson’s reciprocal index (SR) were analyzed with linear ANOVAs (n=5 per treatment at each sampling date) to determine any significant variance in the measured variables that can be attributed to the time of sampling, environmental disturbance (heat, high humidity, control), pesticide treatment (three different a.i and their mixture) and dose (1x, 10x) combinations on microbial parameters. Post-hoc analyses were performed with Tukey’s honestly significant difference test (*p* value < 0.05) using the *TukeyHSD()* function from the *stats* package in RStudio.

The data from β-diversity weighted UniFrac were analyzed with PermANOVA (n=5) using the function adonis from the R package vegan to detect significant differences between time, environmental disturbance, pesticide and dose combinations on bacterial community composition. Post-hoc analyses with pairwise comparisons were performed when significant differences were observed (*p* value < 0.05).

To estimate which OTUs significantly differ between our experimental conditions, we used a generalized linear mixed model. This model accounts for the non-normal distribution of abundance data that typically follow a Poisson distribution, and contains both fixed effects (experimental conditions: time of sampling, environmental disturbances, pesticide treatments and doses) and random effects (sampling effects). Multiple pairwise comparisons were performed with the *emmeans()* function of the *emmeans* package with the Bonferroni correction for p-value adjustment.

## RESULTS

We assessed here the impact of environmental disturbances (heat or high humidity) and a.i exposure (three different a.i and their mixture) on several endpoints: *i*) abundance of 16S, ITS and 18S communities, *ii*) structure and composition of the soil bacterial community (α– and β-diversity) and *iii*) N-cycle functional guilds, by targeting the nitrification step by different means (NH_4_^+^ and NO_3_^−^ pools, AOA, AOB, ComaA and ComaB abundances).

### Effect of environmental disturbances on microbial endpoints

We performed a Principal Component Analysis (PCA) to decipher the contribution of each measured endpoint to the observed response of soil microcosms to environmental disturbances as compared to the control at each time of sampling (T30 and T60, Fig.2 panel A and panel B, respectively). Most of the variance was explained by the two first axes (respectively ∼75 % and ∼65 % at T30 and T60) and separated the heat-disturbed microcosms from the high humidity and the control ones (Fig. 2). At both timepoints, this was mostly explained by α-diversity indices and NO_3_^−^ pool (Table S1).

**Figure 2.**
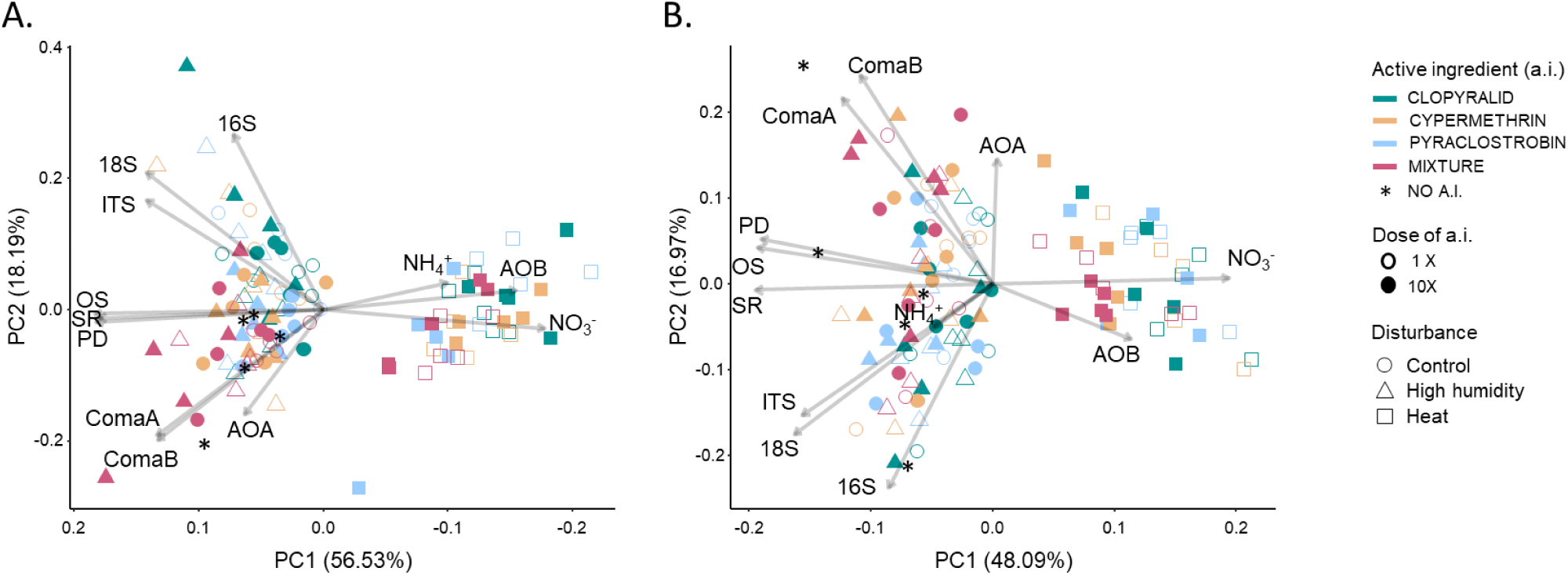
Principal Component Analysis (PCA) of the microbial endpoints (abundance of microbial communities 16S, 18S and ITS; bacterial diversity indices OS, PD and SR; N-cycle functional guilds: abundance of AOA, AOB, ComaA and comaB and abundance of nitrate (NO_3_^−^) and ammonium (NH_4_^+^) pools) measured at T30 (panel A) and T60 (panel B). PC1 and PC2 axes represent the two main principal components capturing the maximum variability in the dataset. Arrows indicate the direction and magnitude of each endpoint’s contribution to the first and second principal components. Each soil microcosm is depicted by a symbol whose shape, color and fill respectively indicate the disturbance, the active ingredient (a.i) and the a.i. doses that were applied.

Focusing on disturbed and control microcosms irrespectively of the applied a.i, we observed significant differences between heat-disturbed microcosms and the others (Table 1). The abundances of 16S, ITS and 18S communities, as well as the diversity of the bacterial community (OS, PD and SR) were significantly lower in heat-disturbed microcosms (ANOVAs followed by Tukey HSD tests; p < 0.05) as compared to the control. When focusing on N-cycle, we also observed a significant decrease in the abundance of the ComaA and ComaB guilds and a significant increase in NH_4_^+^ (only at T30) and NO_3_^−^ pools, as well as in the abundance of AOB in heat-disturbed microcosms. For most of the endpoints, the high humidity disturbance did not induce changes as compared to control microcosms, except for the diversity level of the bacterial community at T30, and the abundance of the protist community (18S).

**Table 1.**
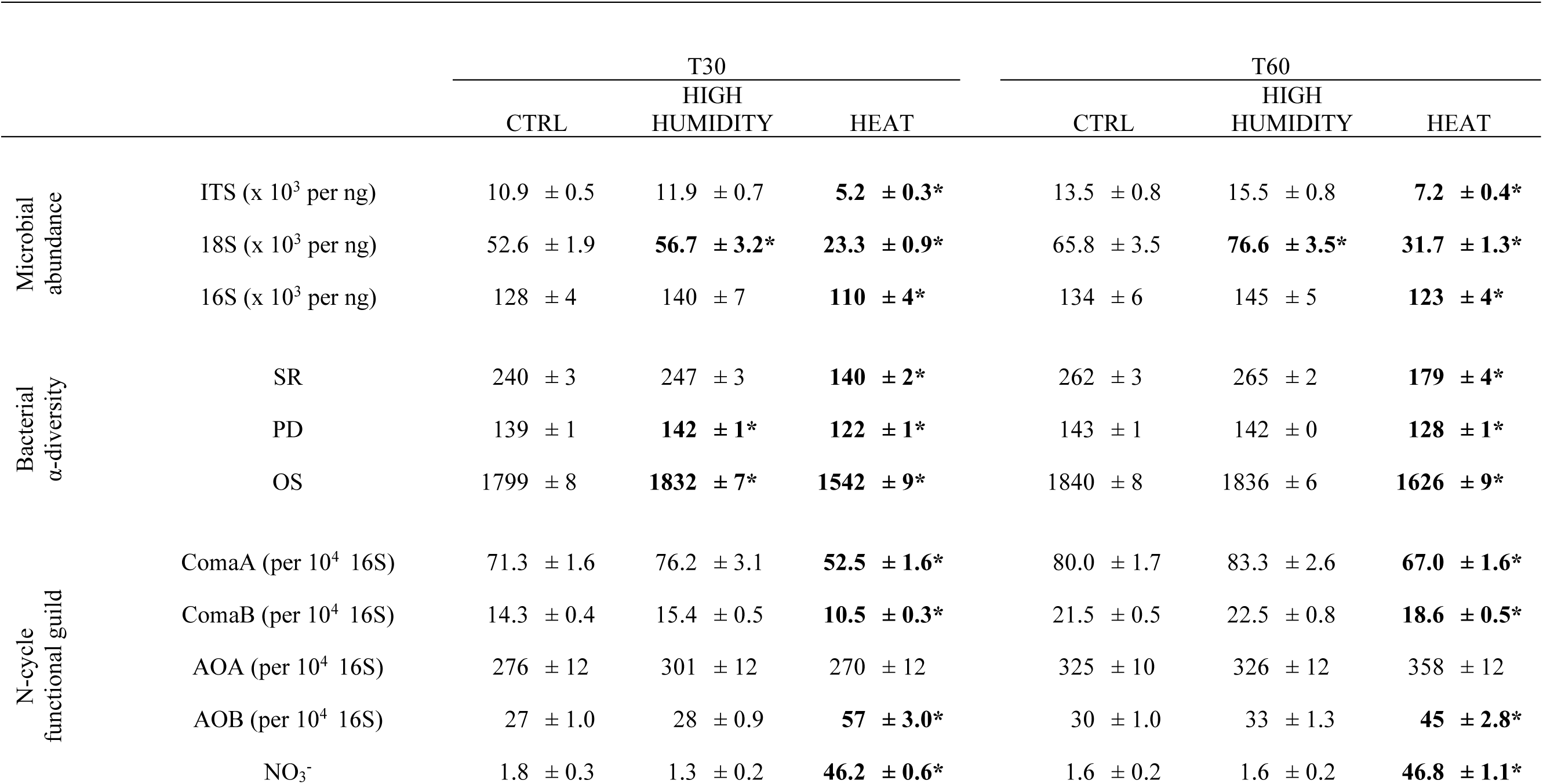
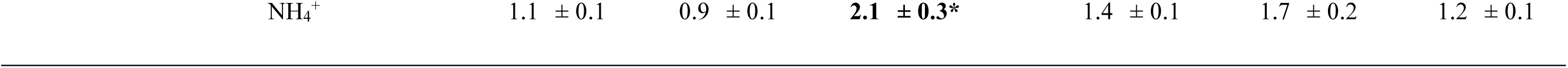
Microbial endpoints (abundance of microbial communities 16S, 18S and ITS; bacterial diversity indices OS, PD and SR; N-cycle functional guilds: abundance of AOA, AOB, ComaA and comaB and abundance of nitrate (NO_3_^−^) and ammonium (NH_4_^+^) pools) measured at T30 and T60 on disturbed (HEAT, HIGH HUMIDITY) and control (CTRL) soil microcosms. For each endpoint, the mean ± the standard error across all pesticide treated microcosms (n= 40 for each disturbance treatment and control) measured at each time point and for each soil disturbance is indicated. For a given time, an asterisk (*) indicates that the endpoint is significantly different from the one measured in the undisturbed soil (ANOVA followed by a Tukey HSD test; p < 0.05). SR: Simpson Reciprocal; PD: PD Whole Tree; OS: Observed Species.

We also performed a Principal Coordinates Analysis (PCoA) of the weighted UniFrac distance matrix to examine bacterial compositional changes in response to environmental disturbances applied to soil microcosms (Fig. 3). In line with results presented above, heat-disturbed microcosms were significantly different from the other two disturbance treatments (high humidity and control) at both timepoints (PermANOVAs; p < 0.05). In addition, the high-humidity-disturbed microcosms were significantly slightly different from control microcosms at T30. Overall, among the 336 dominant OTUs (defined as those having abundance > 0.1 %), ∼80% showed at both T30 and T60, significant differences in their abundance between heat-disturbed and control microcosms, while less than 5% displayed significant differences between high-humidity-disturbed and control ones (Fig. S1).

**Figure 3.**
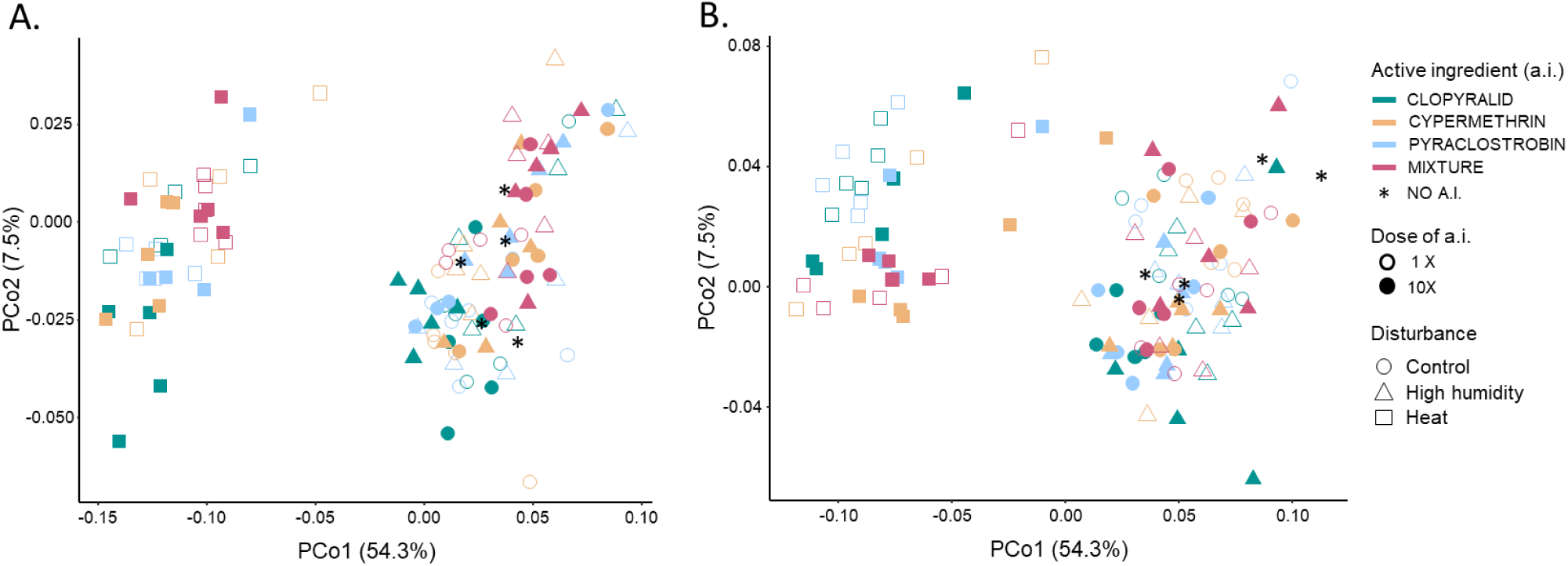
Principal Coordinate Analysis (PCoA) of the bacterial community structure in the different soil microcosms based on the weighted UniFrac distance metric. The two main principal coordinates PCo1 and PCo2 captured respectively 54.3 % and 7.5 % of the observed variability between bacterial communities. For T30 (panel A) and T60 (panel B), each sample is represented by a symbol whose shape, color and fill respectively indicate the disturbance, the active ingredient and the a.i. dose that were applied.

### Evaluating the impact of active ingredients on microbial endpoints

We then focused on the results obtained for the disturbance control microcosms exposed to a.i applied alone (clopyralid, *i. e.* CLO; cypermethrin *i. e.* CYP; pyraclostrobin, *i. e.* PYR) or as mixture (*i. e.* MIX) to evaluate their ecotoxicological impacts on measured endpoints. When considering all the endpoints together in a PCA (Fig. 4, panels A and B), we did not observe a strong clustering of the samples, neither according to the a.i or mixture of the three a.i, nor to the dose of applications (1x or 10x) at both timepoints (T30 and T60). However, when considering the individual endpoints, we detected a significant effect of a.i treatments on NH_4_^+^ and NO_3_^−^ pools, as well as on the abundance of AOA, ComaA and ComaB (even though no significant differences were detected for those groups in pairwise comparisons with the untreated control microcosms) and to a smaller extent on the abundance of 16S, 18S and ITS (ANOVAs, *Pesticide_Dose* and/or *Pesticide_Dose-by-Time* effects p < 0.05). At T30, the NO_3_^−^ pool was the most sensitive endpoint, with significantly higher levels detected in the CLO1x, CLO10x, CYP1x, PYR1x and MIX1x treated soil microcosms as compared to the untreated controls (Table 2). The NH_4_^+^ pool was significantly higher only in the CLO1x and MIX1x treated microcosms as compared to the untreated control. At T60, the 16S abundance was the sole endpoint that was significantly decreased in response to exposure to CYP1x and PYR1x.

**Figure 4.**
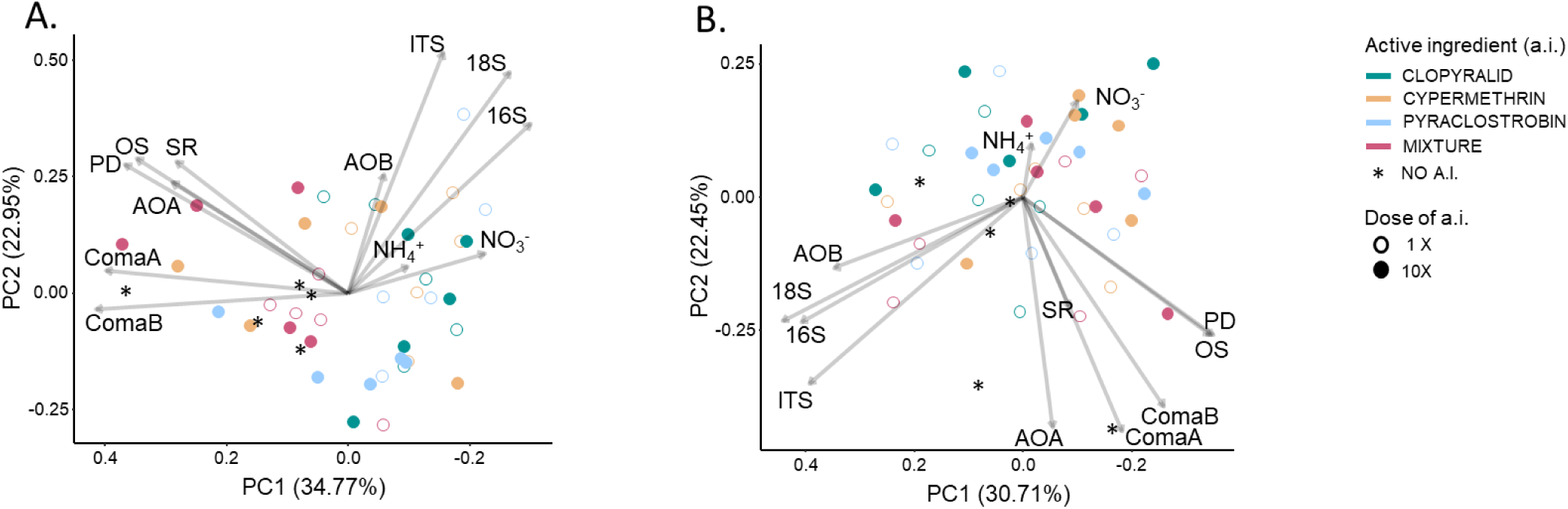
Principal Component Analysis (PCA) performed on the microbial endpoints (abundance of microbial communities 16S, 18S and ITS; bacterial diversity indices OS, PD and SR; N-cycle functional guilds: abundance of AOA, AOB, ComaA and comaB and abundance of nitrate (NO_3_^−^) and ammonium (NH_4_^+^) pools) measured in the undisturbed microcosms at T30 (panel A) and T60 (panel B). The PC1 and PC2 axes represent the two main principal components capturing the maximum variability in the dataset. Arrows indicate the direction and magnitude of each endpoint’s contribution to the first and second principal components. Each soil microcosm is depicted by a circle whose color denotes the active ingredient that it received. Empty circles represent samples having received the a.i. at the agronomical dose while the full circle represent the one who received 10 times the agronomical dose.

**Table 2.**
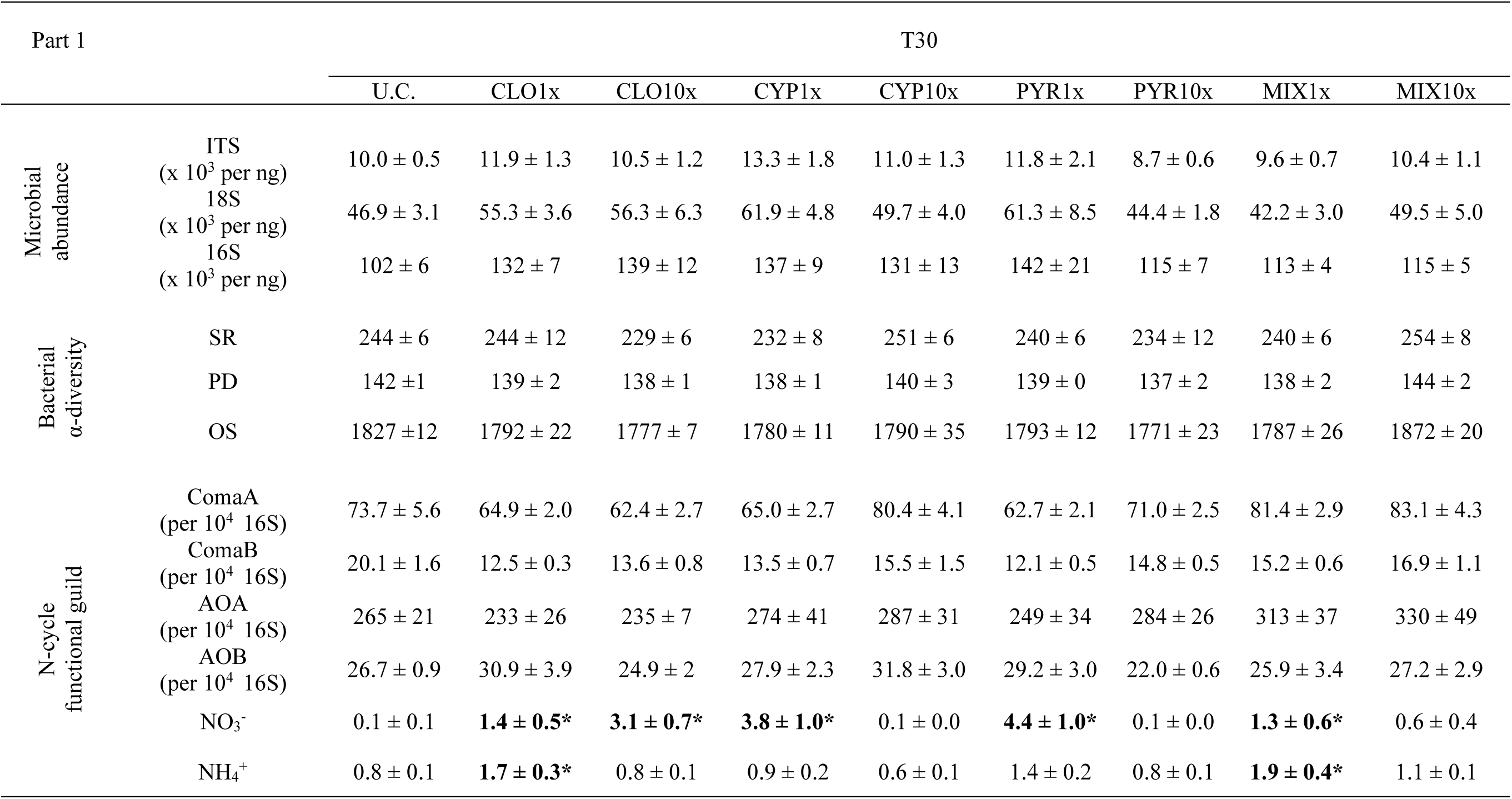
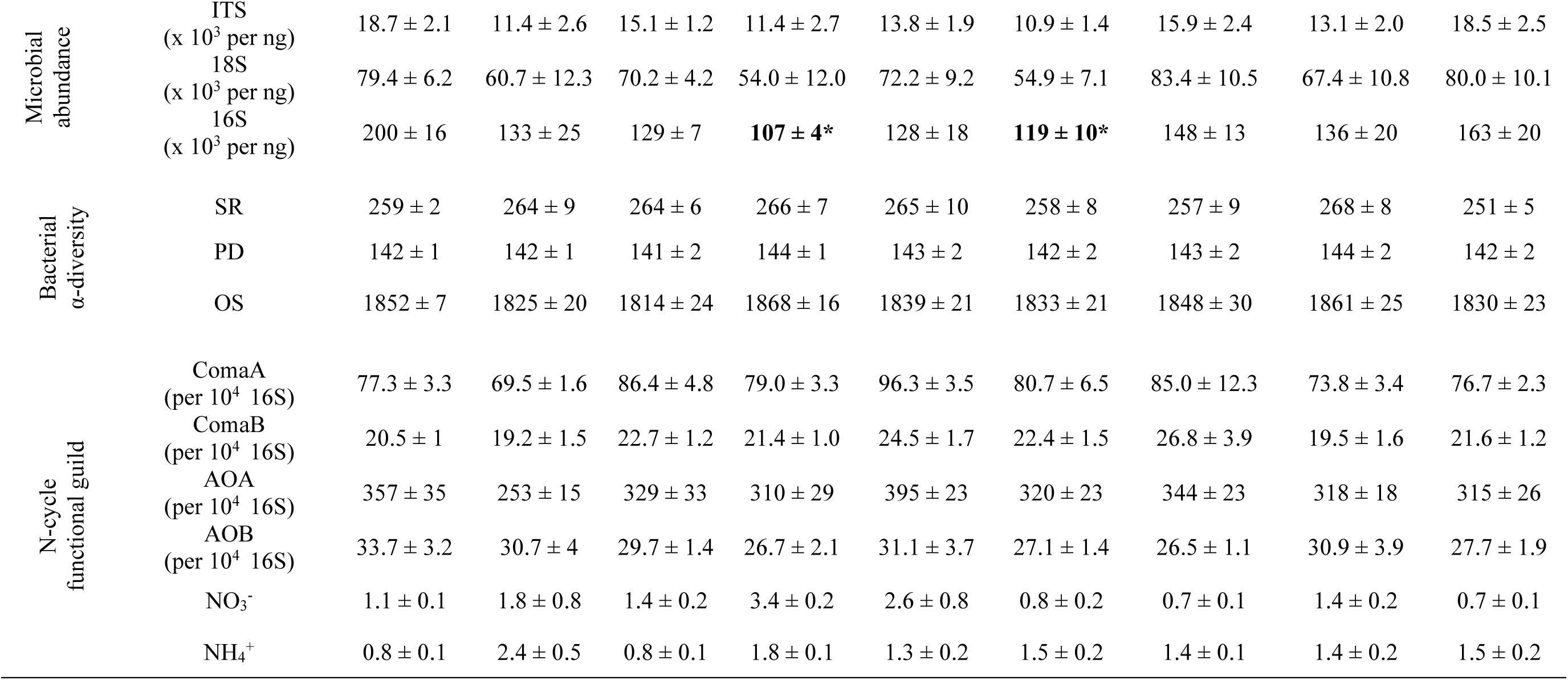
Microbial endpoints (abundance of microbial communities 16S, 18S and ITS; bacterial diversity indices OS, PD and SR; N-cycle functional guilds: abundance of AOA, AOB, ComaA and comaB and abundance of nitrate (NO_3_^−^) and ammonium (NH_4_^+^) pools) measured at T30 and T60 on the undisturbed treated and untreated control (u.c.) soil microcosms. For each endpoint, the mean ± the standard error across all control treated microcosms and u.c. (n=5 for each active ingredient-dose and u.c.) measured at each time point and for each active ingredient is indicated. For a given time, an asterisk (*) indicates that the endpoint is significantly different from the one measured in the u.c. soil (ANOVA followed by a Tukey HSD test; p < 0.05). SR: Simpson Reciprocal; PD: PD Whole Tree; OS: Observed Species.

PCoA on the Weighted UniFrac distance matrix (Fig. 5) showed that in the absence of heat or humidity disturbance the bacterial community structure was not changed by a.i exposure as compared to the untreated control. We noticed that the structure of the bacterial communities of the microcosms exposed to CLO10x were significantly different from those exposed to CYP10x (Pairwise PermANOVAs, p < 0.05). When looking at the dominant OTUs, at both sampling times, we found only 12 OTUs that showed significant differences in relative abundance in a.i treated microcosms versus the untreated control ones (4 Acidobacterias, 3 Actinobacterias, 2 Verrucomicrobias, 2 Proteobacteria and 1 Chloroflexi). Among those, only four were consistently and significantly impacted at both timepoints: they were observed only in CLO-treated microcosms (three in CLO10x and one in CLO1x) and three of them are related to uncultured members of the Acidobacteria phylum.

**Figure 5.**
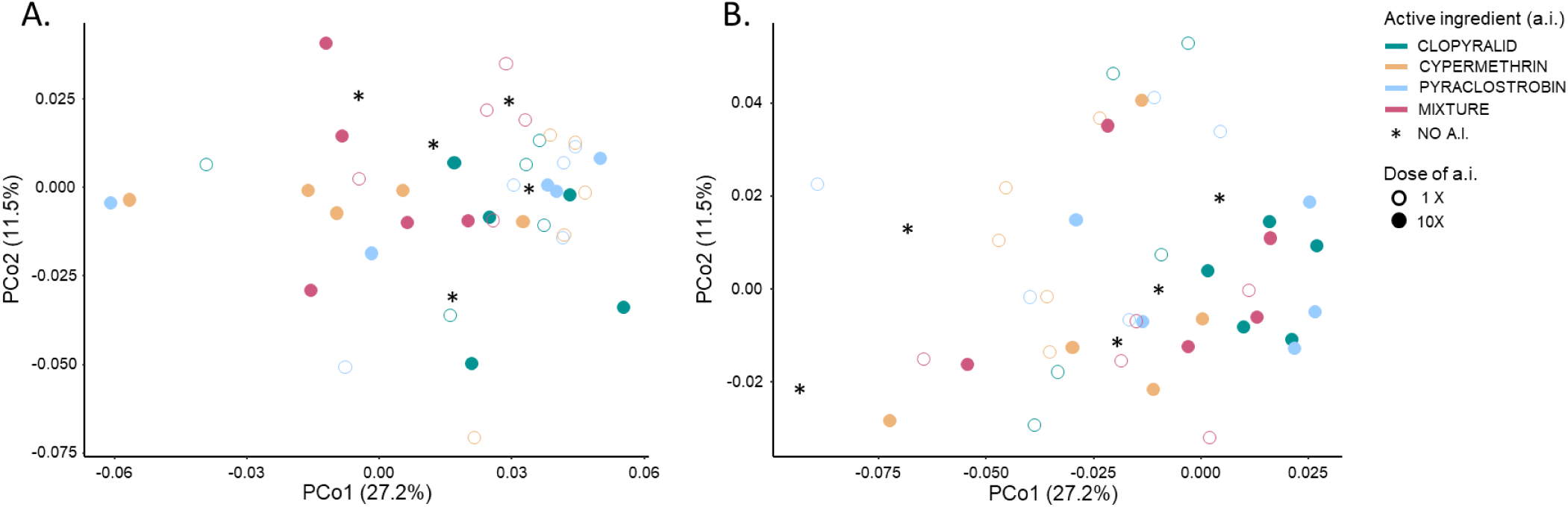
Principal Coordinate Analysis (PCoA) of the bacterial community structure in the undisturbed microcosms based on weighted UniFrac distance metric. The two main principal coordinates PCo1 and PCo2 captured respectively 27.2% and 11.5% of the variability observed between the bacterial communities. For T30 (panel A) and T60 (panel B), each sample is represented by a symbol whose color and fill respectively indicate the active ingredient and its dose applied in soil microcosms.

### Estimating compounded effects of environmental disturbances and pesticides inputs on microbial endpoints

A compounded effect in our ANOVA model would manifest with a significant two-way *‘Disturbance-by-Pesticide_Dose’* or three-way *‘Disturbance-by-Pesticide_Dose-by-TimeOfSampling’* effect, indicating that the effect of a given *‘Pesticide_Dose’* at a given *‘TimeOfSampling’* is conditioned by the *‘Disturbance’*. The AOB abundance and the OS of the bacterial community were the only two endpoints showing a significant three-way interaction (Table 3). However, none of the pairwise comparisons were significant for OS, while the only significant, but slight, difference observed for AOB was between T30-Heat-PYR10x (38 ± 8 copies per 10^4^ copies 16S rDNA) and T30-Heat-CLO10x (75 ± 2 copies per 10^4^ copies 16S rDNA) samples, indicating that the heat disturbance conditioned the relative effects of PYR and CLO at the 10x dose. The two-way interaction was only significant for the abundance of 16S, and NH_4_^+^ and NO_3_^−^ pools. More specifically, CYP1x-treated microcosms showed significantly higher levels of NO_3_^−^ compared to PYR1x, PYR10x and CYP10x-treated microcosms, but only in the treated controls, while PYR10x-treated microcosms displayed significantly higher levels of NH_4_^+^ than PYR1x, CYP1x, CLO1x and MIX10x-treated ones, but only for the heat-disturbed microcosms. Overall, compounded effects are relatively sparse, idiosyncratic and of small scale which is in line with the relatively small ecotoxicological impact detected for the chosen pesticides in undisturbed environmental conditions.

**Table 3.** Analysis of variance (ANOVA) results assessing the impact of time of sampling, the disturbance, and the applied pesticide (nature/dose) on various microbial endpoints (16S: bacterial abundance; OS: Observed Species; AOB: Ammonia Oxydizing Bacteria abundance) and nitrate (NO. _3_**^−^) and ammonium pools (NH**_4_**^+^).** Only the endpoints for which a significant 2-way *‘Disturbance-by-Pesticide_Dose’* or 3-way *‘Disturbance-by-Pesticide_Dose-by-TimeOfSampling’* interactions are presented (p < 0.05).

|  |  | 16S |  |  | OS |  |  | AOB |  |  | NO<sub>3</sub><sup>-</sup> |  |  | NH<sub>4</sub><sup>+</sup> |  |  |
| --- | --- | --- | --- | --- | --- | --- | --- | --- | --- | --- | --- | --- | --- | --- | --- | --- |
|  | Df | Mean<br>Sq | F<br>value | Pr(>F) | Mean<br>Sq<br>x10<sup>3</sup> | F<br>value | Pr(>F) | Mean<br>Sq | F<br>value | Pr(>F) | Mean<br>Sq | F<br>value | Pr(>F) | Mean<br>Sq | F<br>value | Pr(>F) |
| *TimeOfSampling* | 1 | 0.04 | 4.3 | 0.040 | 114 | 60 | < 0.001 | 0.00 | 0 | 0.985 | 1.90 | 4.70 | 0.031 | 0.537 | 13.4 | < 0.001 |
| *Disturbance* | 2 | 0.14 | 13.5 | < 0.001 | 1569 | 821 | < 0.001 | 1.31 | 114 | < 0.001 | 1.35 | 3.32 | 0.038 | 0.130 | 3.2 | 0.041 |
| *Pesticide_Dose* | 7 | 0.02 | 2.2 | 0.037 | 10 | 5 | < 0.001 | 0.01 | 1 | 0.554 | 0.55 | 1.36 | 0.224 | 0.151 | 3.8 | 0.001 |
| *Disturbance-by-TimeOfSampling* | 2 | 0.01 | 0.8 | 0.466 | 32 | 17 | < 0.001 | 0.15 | 13 | < 0.001 | 0.06 | 0.16 | 0.852 | 0.964 | 24.1 | < 0.001 |
| *Pesticide_Dose-by-TimeOfSampling* | 7 | 0.01 | 1.1 | 0.373 | 3 | 2 | 0.089 | 0.02 | 2 | 0.053 | 0.81 | 1.99 | 0.059 | 0.145 | 3.6 | 0.001 |
| *Disturbance-by-Pesticide_Dose* | 14 | 0.02 | 1.9 | 0.030 | 3 | 1 | 0.192 | 0.01 | 1 | 0.263 | 1.37 | 3.40 | < 0.001 | 0.133 | 3.3 | < 0.001 |
| *Disturbance-by-Pesticide_Dose-by-TimeOfSampling* | 14 | 0.01 | 0.7 | 0.781 | 4 | 2 | 0.013 | 0.03 | 2 | 0.007 | 0.68 | 1.69 | 0.061 | 0.064 | 1.6 | 0.084 |

## DISCUSSION

The present study aimed to evaluate the effects of elevated temperature (heat) and heavy rainfall (high humidity), two global change related environmental disturbances, and of three commonly used a.i., CLO (herbicide), CYP (insecticide), PYR (fungicide) and their mixture on soil microbial community abundance, structure, composition and functioning. We used qPCR techniques to detect any effect on the abundance of some important microbial guilds, and Illumina next generation sequencing of the 16S rDNA amplicons to study the effects on the bacterial community composition.

Most of the measured microbial endpoints were affected by the heat stress, with significant effects at T30 (short term). The α-diversity indices and important N-cycle-related microbial guilds were the endpoints mostly impacted by the heat disturbance. We found decreasing α-diversity indices and accumulation of NO_3-_, which is in line with former studies in which the disruption of the N-cycle was described as a consequence of a lower functional redundancy due to community shifts (Calderón et al., 2018). In another research, the heat-disturbed microbial communities never recovered, and were always different from the control, independently from the soil disturbance history (Jurburg et al., 2017a). This response is in line with our observations, where the effect of heat stress persisted all along the experiment. We also measured a higher AOB abundance in the heat disturbed samples compared to the control, which suggests a higher tolerance to heat of the AOB community compared to other members of N microbial guilds. This finding is supported by other studies, in which AOB were more responsive to soil warming than AOA (Szukics et al., 2010; Wang, 2021). We observed that ComaA and ComaB abundances were negatively impacted by the heat stress: not much is known about the effect of heat on these two microbial guilds, but a recent research suggests that commamox are favoured at low temperature in wastewater treatment plants (Gonzalez-Martinez et al., 2016). We can therefore conclude that the N-cycle was impacted by heat stress at several levels, and exhibited a limited recovery, *i. e.* low resilience, at the time scale of the experiment.

The effect of pesticides on soil microbial communities has been studied for decades (Bollen, 1961). The microbial response to pesticide application is variable, and depends on many factors such as the pesticidal mode of action, the experimental set up and the studied endpoints (Jacobsen and Hjelmsø, 2014; Puglisi, 2012). Indeed, the few responses observed in our experiment are not consistent across the various endpoints analysed. Most are related to the N-cycle and exhibit a transient effect, *i. e.* recovery in the longer term. Some functional guilds, e.g. AOA and AOB, have been described to be sensitive to pesticide application (Karas et al., 2018). To our best knowledge, no studies have explored the influence of pesticides on comammox. Even though we could not detect a strong effect, our findings indicate that the application of technical a.i might affect these N-cycle microbial guilds. The exposure of soil to the a.i, and particularly to the herbicide CLO, led to a significant increase of the NO_3_^−^ and NH_4_^+^ pools which is supported by previous observations reporting the disruption of the N-cycle in response to pesticide application (Brochado et al., 2023; Damin and Trivelin, 2011; Hernández et al., 2011). Interestingly, one could observe that exposure to a.i had significant effects on the above mentioned endpoints already at agronomical dose, contrary to what is reported in the existing literature where effects are mostly detected at higher doses (Crouzet et al., 2010; Puglisi, 2012; Romdhane et al., 2019). The application of a.i alone or in a mixture had no strong impact on microbial community composition. This might be due to the large variability observed between biological replicates within given experimental conditions and to the limited effect of the pesticide. Among all the OTUs, just three affiliated to the Acidobacteria were responsive to one pesticide (CLO). This phylum is very abundant and ubiquitous in many ecosystems, and takes part to various metabolic pathways like carbon and nitrogen cycle (Kalam et al., 2020). Previous studies described this phyla to be a good biological indicator of land-use change from forest to farmland because of its sensitivity to various toxic metals, potentially deriving from metal containing pesticides (Kim et al., 2021).

The three a.i used in our study are considered from EFSA as showing low toxicity towards soil living organisms (European Food Safety Authority (EFSA) et al., 2018a, 2018b, 2018c), but nothing is known about the effect of these three compounds applied in combination. Contrary to our expectation, the exposure of soil microcosms to the mixture of these three a.i at the agronomic application rate and at 10-fold this rate did not affect the measured endpoints differently than exposure to the respective single a.i. Studies on the effect of pesticide mixtures on the soil microbial community are scarce (Baćmaga et al., 2015; Joly et al., 2015, 2012; Schuster and Schröder, 1990), and the responses are variable. Based on knowledge for other endpoints (Cedergreen 2014), the probability of detecting synergistic effects is low. Hence, we can conclude that the present experiment confirms the findings of Cedergreen (2014) as no exceptionally stronger effect was found in the mixture treatments (pointing at a potential synergistic interaction) compared to the single a.i. treatments.

A compounded disturbance is the phenomenon in which ecosystems are subjected to multiple disturbances occurring over time. The effect of the initial disturbance may have legacy effect and consequently alter the ability of the microbial community to cope with the following ones. The responses to the following stresses are variable (Vinebrooke et al., 2004): *i. e.* the exposure to multiple perturbations of the same nature might lead to microbial selection and high community resilience to that specific stress (Calderón et al., 2018). However, if the subsequent perturbation is of a different nature, the disturbed community might either become more resistant to the following stress because of community priming (Rillig et al., 2015), or more sensitive (Calderón et al., 2018) because of negative species co-tolerance (Vinebrooke et al., 2004). In our study, the application of different a.i to a previously environmentally disturbed community had no apparent further impact on the microbial communities. Responses to a.i application were few and transient, but it is worth mentioning that the heat stress seemed to condition the response to the different tested a.i. This was the case of AOB or NH_4_^+^, especially to PYR treatment. According to the literature, abiotic disturbances, such as extreme temperatures show a great potential into shaping and modifying the microbial community (Bardgett and Caruso, 2020; Castro et al., 2010; Islam et al., 2020) as compared to pesticides where the effects are often variable and confined to certain endpoints.

Overall, the temperature disturbance had a major and persisting impact on microbial communities. On the opposite, the transient increase in soil humidity was without effects. Similarly, the effects of the tested a.i and their mixture on the control community were only slight and transient, and seem to impact mainly some N-cycle endpoints. We did not find any strong and persisting effect on microbial endpoints of the compounded disturbances. Future studies should be conducted in different soils with various physico-chemical properties to generalize our conclusions. Also, consideration of denitrifiers and N_2_O emitters would be of interest to give a more comprehensive understanding of the compounded effect of environmental disturbances and pesticides on N-cycle in soils.

## Acknowledgments

The project leading to this publication, as well as Camilla Drocco’s PhD, have received funding from the European Union’s Horizon 2020 research and innovation programme under the Marie Skłodowska-Curie grant agreement No 956496

## Notes

### Competing Interest Statement

The authors have declared no competing interest.

### Summary of Updates

This version has been Peer-reviewed & recommended by PCI Ecotox Env Chem.

## REFERENCES

1. Aislabie, J., Deslippe, J.R., 2013. Soil microbes and their contribution to soil services, in: Ecosystem Services in New Zealand – Conditions and Trends. Dymond, J.R. Ed., Manaaki Whenua Press, Lincoln, NZ, pp. 143.161

2. Altschul, S.F., Gish, W., Miller, W., Myers, E.W., Lipman, D.J., 1990. Basic Local Alignment Search Tool. J. Mol. Biol. 215(3), 403–410. 10.1016/S0022-2836(05)80360-2

3. Baćmaga, M., Borowik, A., Kucharski, J., Tomkiel, M., Wyszkowska, J., 2015. Microbial and enzymatic activity of soil contaminated with a mixture of diflufenican + mesosulfuron-methyl + iodosulfuron-methyl-sodium. Environ. Sci. Pollut. Res. 22, 643–656. 10.1007/s11356-014-3395-5

4. Bardgett, R.D., Caruso, T., 2020. Soil microbial community responses to climate extremes: resistance, resilience and transitions to alternative states. Philos. Trans. R. Soc. B. 375, 20190112. 10.1098/rstb.2019.0112

5. Barnard, R.L., Osborne, C.A., Firestone, M.K., 2013. Responses of soil bacterial and fungal communities to extreme desiccation and rewetting. ISME J. 7, 2229–2241. 10.1038/ismej.2013.104

6. Berry, D., Mahfoudh, K.B., Wagner, M., Loy, A., 2011. Barcoded Primers Used in Multiplex Amplicon Pyrosequencing Bias Amplification. Appl. Environ. Microbiol. 77, 7846–7849. 10.1128/AEM.05220-11

7. Bollen, W.B., 1961. Interactions Between Pesticides and Soil Microorganisms. Annu. Rev. Microbiol. 15, 69–92. 10.1146/annurev.mi.15.100161.000441

8. Brochado, M.G. d. S., Silva, L.B.X. d., Lima, A. d. C., Guidi, Y.M., Mendes, K.F., 2023. Herbicides versus Nitrogen Cycle: Assessing the Trade-Offs for Soil Integrity and Crop Yield—An In-Depth Systematic Review. Nitrogen 4, 296–310. 10.3390/nitrogen4030022

9. Bru, D., Ramette, A., Saby, N., Dequiedt, S., Ranjard, L., Jolivet, C., Arrouays, D., Philippot, L., 2011. Determinants of the distribution of nitrogen-cycling microbial communities at the landscape scale. ISME J. 5, 532–542. 10.1038/ismej.2010.130

10. Calderón, K., Philippot, L., Bizouard, F., Breuil, M.-C., Bru, D., Spor, A., 2018. Compounded Disturbance Chronology Modulates the Resilience of Soil Microbial Communities and N-Cycle Related Functions. Front. Microbiol. 9, 2721. doi: 10.3389/fmicb.2018.02721

11. Caporaso, J.G., Kuczynski, J., Stombaugh, J., Bittinger, K., Bushman, F.D., Costello, E.K., Fierer, N., Peña, A.G., Goodrich, J.K., Gordon, J.I., Huttley, G.A., Kelley, S.T., Knights, D., Koenig J.E., Ley, R.E., Lozupone, C.A., McDonald, D., Muegge, B.D., Pirrung, M., Reeder, J., Sevinsky, J.R., Turnbaugh, P.J., Walters, W.A., Widmann, J., Yatsunenko, T., Zaneveld, J., Knight, R., 2010. QIIME allows analysis of high-throughput community sequencing data. Nat Method 7(5), 335–336. doi:10.1038/nmeth.f.303

12. Castro, H.F., Classen, A.T., Austin, E.E., Norby, R.J., Schadt, C.W., 2010. Soil Microbial Community Responses to Multiple Experimental Climate Change Drivers. Appl. Environ. Microbiol. 76(4), 999– 1007. doi:10.1128/AEM.02874-09

13. Cedergreen, N., 2014. Quantifying Synergy: A Systematic Review of Mixture Toxicity Studies within Environmental Toxicology. PLoS ONE 9, e96580. 10.1371/journal.pone.0096580

14. Crouzet, O., Batisson, I., Besse-Hoggan, P., Bonnemoy, F., Bardot, C., Poly, F., Bohatier, J., Mallet, C., 2010. Response of soil microbial communities to the herbicide mesotrione: A dose-effect microcosm approach. Soil Biol. Biochem. 42, 193–202. 10.1016/j.soilbio.2009.10.016

15. Damin, V., Trivelin, P.C.O., 2011. Herbicides Effect on Nitrogen Cycling in Agroecosystems, in: Herbicides and Environment. Kortekamp, A., Ed. IntechOpen, Rijeka, pp. 107–124.

16. Dominati, E., Patterson, M., Mackay, A., 2010. A framework for classifying and quantifying the natural capital and ecosystem services of soils. Ecol. Econ. 69, 1858–1868. 10.1016/j.ecolecon.2010.05.002

17. European Food Safety Authority (EFSA), Arena, M., Auteri, D., Barmaz, S., Brancato, A., Brocca, D., Bura, L., Carrasco Cabrera, L., Chiusolo, A., Civitella, C., Court Marques, D., Crivellente, F., Ctverackova, L., De Lentdecker, C., Egsmose, M., Erdos, Z., Fait, G., Ferreira, L., Greco, L., Ippolito, A., Istace, F., Jarrah, S., Kardassi, D., Leuschner, R., Lostia, A., Lythgo, C., Magrans, J.O., Medina, P., Mineo, D., Miron, I., Molnar, T., Padovani, L., Parra Morte, J.M., Pedersen, R., Reich, H., Sacchi, A., Santos, M., Serafimova, R., Sharp, R., Stanek, A., Streissl, F., Sturma, J., Szentes, C., Tarazona, J., Terron, A., Theobald, A., Vagenende, B., Van Dijk, J., Villamar-Bouza, L., 2018a. Peer review of the pesticide risk assessment of the active substance clopyralid. EFSA J. 16(8), 5389. 10.2903/j.efsa.2018.5389

18. European Food Safety Authority (EFSA), Arena, M., Auteri, D., Barmaz, S., Brancato, A., Brocca, D., Bura, L., Carrasco Cabrera, L., Chiusolo, A., Civitella, C., Court Marques, D., Crivellente, F., Ctverackova, L., De Lentdecker, C., Egsmose, M., Erdos, Z., Fait, G., Ferreira, L., Greco, L., Ippolito, A., Istace, F., Jarrah, S., Kardassi, D., Leuschner, R., Lostia, A., Lythgo, C., Magrans, J.O., Medina, P., Mineo, D., Miron, I., Molnar, T., Padovani, L., Parra Morte, J.M., Pedersen, R., Reich, H., Sacchi, A., Santos, M., Serafimova, R., Sharp, R., Stanek, A., Streissl, F., Sturma, J., Szentes, C., Tarazona, J., Terron, A., Theobald, A., Vagenende, B., Van Dijk, J., Villamar-Bouza, L., 2018b. Peer review of the pesticide risk assessment of the active substance cypermethrin. EFSA J. 16(8), 5402. doi: 10.2903/j.efsa.2018.5402

19. European Food Safety Authority, 2018c. Public consultation on the active substance pyraclostrobin. https://www.efsa.europa.eu/en/consultations/call/180710

20. Faith, D.P., 1992. Conservation evaluation and phylogenetic diversity. Biol. Conserv. 61, 1–10. 10.1016/0006-3207(92)91201-3

21. Food and Agriculture Organization of the United Nations (2023). FAOSTAT, Pesticide Use Database. Accessed: December 2023. https://www.fao.org/faostat/en/#data/RP

22. Franco-Andreu, L., Gómez, I., Parrado, J., García, C., Hernández, T., Tejada, M., 2016. Behavior of two pesticides in a soil subjected to severe drought. Effects on soil biology. Appl. Soil Ecol. 105, 17–24. 10.1016/j.apsoil.2016.04.001

23. Frey, S.D., Drijber, R., Smith, H., Melillo, J., 2008. Microbial biomass, functional capacity, and community structure after 12 years of soil warming. Soil Biol. Biochem. 40, 2904–2907. 10.1016/j.soilbio.2008.07.020

24. Geissen, V., Silva, V., Lwanga, E.H., Beriot, N., Oostindie, K., Bin, Z., Pyne, E., Busink, S., Zomer, P., Mol, H., Ritsema, C.J., 2021. Cocktails of pesticide residues in conventional and organic farming systems in Europe – Legacy of the past and turning point for the future. Environ. Pollut. 278, 116827. 10.1016/j.envpol.2021.116827

25. Gonzalez-Martinez, A., Rodriguez-Sanchez, A., van Loosdrecht, M.C.M., Gonzalez-Lopez, J., Vahala, R., 2016. Detection of comammox bacteria in full-scale wastewater treatment bioreactors using tag-454-pyrosequencing. Environ. Sci. Pollut. Res. 23, 25501–25511. 10.1007/s11356-016-7914-4

26. Hernández, M., Jia, Z., Conrad, R., Seeger, M., 2011. Simazine application inhibits nitrification and changes the ammonia-oxidizing bacterial communities in a fertilized agricultural soil. FEMS Microbiol. Ecol. 78, 511–519. 10.1111/j.1574-6941.2011.01180.x

27. Islam, W., Noman, A., Naveed, H., Huang, Z., Chen, H.Y.H., 2020. Role of environmental factors in shaping the soil microbiome. Env. Sci Pollut Res 27, 41225–41247. 10.1007/s11356-020-10471-2

28. Jacobsen, C.S., Hjelmsø, M.H., 2014. Agricultural soils, pesticides and microbial diversity. Curr. Opin. Biotechnol. 27, 15–20. 10.1016/j.copbio.2013.09.003

29. Joly, P., Besse-Hoggan, P., Bonnemoy, F., Batisson, I., Bohatier, J., Mallet, C., 2012. Impact of Maize Formulated Herbicides Mesotrione and S-Metolachlor, Applied Alone and in Mixture, on Soil Microbial Communities. ISRN Ecol. 2012, 1–9. 10.5402/2012/329898

30. Joly, P., Bonnemoy, F., Besse-Hoggan, P., Perrière, F., Crouzet, O., Cheviron, N., Mallet, C., 2015. Responses of Limagne “Clay/Organic Matter-Rich” Soil Microbial Communities to Realistic Formulated Herbicide Mixtures, Including S-Metolachlor, Mesotrione, and Nicosulfuron. Water Air Soil Pollut. 226, 413. 10.1007/s11270-015-2683-0

31. Jurburg, S.D., Nunes, I., Brejnrod, A., Jacquiod, S., Priemé, A., Sørensen, S.J., Van Elsas, J.D., Salles, J.F., 2017a. Legacy Effects on the Recovery of Soil Bacterial Communities from Extreme Temperature Perturbation. Front. Microbiol. 8, 1832. 10.3389/fmicb.2017.01832

32. Jurburg, S.D., Nunes, I., Stegen, J.C., Le Roux, X., Priemé, A., Sørensen, S.J., Salles, J.F., 2017b. Autogenic succession and deterministic recovery following disturbance in soil bacterial communities. Sci. Rep. 7, 45691. 10.1038/srep45691

33. Kalam, S., Basu, A., Ahmad, I., Sayyed, R.Z., El-Enshasy, H.A., Dailin, D.J., Suriani, N.L., 2020. Recent Understanding of Soil Acidobacteria and Their Ecological Significance: A Critical Review. Front. Microbiol. 11, 580024. 10.3389/fmicb.2020.580024

34. Karas, P.A., Baguelin, C., Pertile, G., Papadopoulou, E.S., Nikolaki, S., Storck, V., Ferrari, F., Trevisan, M., Ferrarini, A., Fornasier, F., Vasileiadis, S., Tsiamis, G., Martin-Laurent, F., Karpouzas, D.G., 2018. Assessment of the impact of three pesticides on microbial dynamics and functions in a lab-to-field experimental approach. Sci. Total Environ. 637–638, 636–646. 10.1016/j.scitotenv.2018.05.073

35. Kim, H.-S., Lee, S.-H., Jo, H.Y., Finneran, K.T., Kwon, M.J., 2021. Diversity and composition of soil Acidobacteria and Proteobacteria communities as a bacterial indicator of past land-use change from forest to farmland. Sci. Total Environ. 797, 148944. 10.1016/j.scitotenv.2021.148944

36. Kluyver, T., Ragan-Kelley, B., Pérez, F., Granger, B., Bussonnier, M., Frederic, J., Kelley, K., Hamrick, J., Grout, J., Corlay, S., Ivanov, P., Avila, D., Abdalla, S., Willing, C., Jupyter Development Team, 2016. Jupyter Notebooks—a publishing format for reproducible computational workflows. IOS Press. 87–90. 10.3233/978-1-61499-649-1-87

37. Koch, H., van Kessel, M.A.H.J., Lücker, S., 2019. Complete nitrification: insights into the ecophysiology of comammox Nitrospira. Appl. Microbiol. Biotechnol. 103, 177–189. 10.1007/s00253-018-9486-3

38. Kumar, V., Singh, S., Upadhyay, N., 2019. Effects of organophosphate pesticides on siderophore producing soils microorganisms. Biocatal. Agric. Biotechnol. 21, 101359. 10.1016/j.bcab.2019.101359

39. Kuypers, M.M.M., Marchant, H.K., Kartal, B., 2018. The microbial nitrogen-cycling network. Nat. Rev. Microbiol. 16, 263–276. 10.1038/nrmicro.2018.9

40. Lehtovirta-Morley, L.E., 2018. Ammonia oxidation: Ecology, physiology, biochemistry and why they must all come together. FEMS Microbiol. Lett. 365, fny058. 10.1093/femsle/fny058

41. Lozupone, C., Knight, R., 2005. UniFrac: a New Phylogenetic Method for Comparing Microbial Communities. Appl. Environ. Microbiol. 71(12), 8228–8235. 10.1128/AEM.71.12.8228-8235.2005

42. Meisner, A., Jacquiod, S., Snoek, B.L., ten Hooven, F.C., van der Putten, W.H., 2018. Drought Legacy Effects on the Composition of Soil Fungal and Prokaryote Communities. Front. Microbiol. 9, 294. 10.3389/fmicb.2018.00294

43. Moebius-Clune, B.N., Moebius-Clune, D.J., Gugino, B.K., Idowu, O.J., Schindelbeck, R.R., Ristow, A.J., van Es, H.M., Thies, J.E., Shayler, H.A., McBride, M.B., Kurtz, K.S.M., Wolfe, D.W., Abawi, G.S., 2016. Comprehensive assessment of soil health: the Cornell framework manual, 3.2. ed. Cornell University, Geneva, NY.

44. Mooshammer, M., Hofhansl, F., Frank, A.H., Wanek, W., Hämmerle, I., Leitner, S., Schnecker, J., Wild, B., Watzka, M., Keiblinger, K.M., Zechmeister-Boltenstern, S., Richter, A., 2017. Decoupling of microbial carbon, nitrogen, and phosphorus cycling in response to extreme temperature events. Sci. Adv. e1602781. 10.1126/sciadv.1602781

45. Muyzer, G., De Waal, E.C., Uitterlinden, A.G., 1993. Profiling of Complex Microbial Populations by Denaturing Gradient Gel Electrophoresis Analysis of Polymerase Chain Reaction-Amplified Genes Coding for 16S rRNA. Appl. Environ. Microbiol. 59(3), 695–700. 10.1128/aem.59.3.695-700.1993

46. Nawrocki, E.P., Eddy, S.R., 2013. Infernal 1.1: 100-fold faster RNA homology searches. Bioinformatics 29(22), 2933–2935. doi:10.1093/bioinformatics/btt509

47. Paine, R.T., Tegner, M.J., Johnson, E.A., 1998. Compounded Perturbations Yield Ecological Surprises. Ecosystems 1, 535–545. 10.1007/s100219900049

48. Pesaro, M., Nicollier, G., Zeyer, J., Widmer, F., 2004. Impact of Soil Drying-Rewetting Stress on Microbial Communities and Activities and on Degradation of Two Crop Protection Products. Appl. Environ. Microbiol. 70, 2577–2587. 10.1128/AEM.70.5.2577-2587.2004

49. Philippot, L., Griffiths, B.S., Langenheder, S., 2021. Microbial Community Resilience across Ecosystems and Multiple Disturbances. Microbiol. Mol. Biol. Rev. 85, e00026–20. 10.1128/MMBR.00026-20

50. Philippot, L., Spor, A., Hénault, C., Bru, D., Bizouard, F., Jones, C.M., Sarr, A., Maron, P.-A., 2013. Loss in microbial diversity affects nitrogen cycling in soil. ISME J. 7, 1609–1619. 10.1038/ismej.2013.34

51. Pjevac, P., Schauberger, C., Poghosyan, L., Herbold, C.W., van Kessel, M.A.H.J., Daebeler, A., Steinberger, M., Jetten, M.S.M., Lücker, S., Wagner, M., Daims, H., 2017. AmoA-Targeted Polymerase Chain Reaction Primers for the Specific Detection and Quantification of Comammox Nitrospira in the Environment. Front. Microbiol. 8, 1508. 10.3389/fmicb.2017.01508

52. Price, M.N., Dehal, P.S., Arkin, A.P., 2009. FastTree: Computing Large Minimum Evolution Trees with Profiles instead of a Distance Matrix. Mol. Biol. Evo. 26(7), 1641–1650. 10.1093/molbev/msp077

53. Puglisi, E., 2012. Response of microbial organisms (aquatic and terrestrial) to pesticides. EFSA Support. Publ. 9. 10.2903/sp.efsa.2012.EN-359

54. Quast, C., Pruesse, E., Yilmaz, P., Gerken, J., Schweer, T., Yarza, P., Peplies, J., Glöckner, F.O., 2013. The SILVA ribosomal RNA gene database project: improved data processing and web-based tools. Nucleic Acids Res. 41, D590–D596. 10.1093/nar/gks1219

55. Rillig, M.C., Rolff, J., Tietjen, B., Wehner, J., Andrade-Linares, D.R., 2015. Community priming—effects of sequential stressors on microbial assemblages. FEMS Microbiol. Ecol. 91, fiv040. 10.1093/femsec/fiv040

56. Rodríguez Eugenio, N., McLaughlin, M.J., Pennock, D.J., 2018. Soil pollution: a hidden reality. FAO, Rome, IT, pp. 142

57. Rognes, T., Flouri, T., Nichols, B., Quince, C., Mahé, F., 2016. VSEARCH: a versatile open source tool for metagenomics. PeerJ 4, e2584. 10.7717/peerj.2584

58. Romdhane, S., Devers-Lamrani, M., Beguet, J., Bertrand, C., Calvayrac, C., Salvia, M.-V., Jrad, A.B., Dayan, F.E., Spor, A., Barthelmebs, L., Martin-Laurent, F., 2019. Assessment of the ecotoxicological impact of natural and synthetic β-triketone herbicides on the diversity and activity of the soil bacterial community using omic approaches. Sci. Total Environ. 651, 241–249. 10.1016/j.scitotenv.2018.09.159

59. Sahrawat, K.L., 2008. Factors Affecting Nitrification in Soils. Commun. Soil Sci. Plant Anal. 39, 1436– 1446. 10.1080/00103620802004235

60. Schuster, E., Schröder, D., 1990. Side-Effects of Sequentially– and Simultaneously-Applied Pesticides on Non-Target Soil Microorganisms: Laboratory Experiments. Soil Biol. Biochem. 22(3), 375–383. 10.1016/0038-0717(90)90116-H

61. Singh, J.S., 2015. Microbes Play Major Roles in the Ecosystem Services. Clim. Change Environ. Sustain. 3, 163–167. 10.5958/2320-642X.2015.00018.6

62. Storck, V., Nikolaki, S., Perruchon, C., Chabanis, C., Sacchi, A., Pertile, G., Baguelin, C., Karas, P.A., Spor, A., Devers-Lamrani, M., Papadopoulou, E.S., Sibourg, O., Malandain, C., Trevisan, M., Ferrari, F., Karpouzas, D.G., Tsiamis, G., Martin-Laurent, F., 2018. Lab to Field Assessment of the Ecotoxicological Impact of Chlorpyrifos, Isoproturon, or Tebuconazole on the Diversity and Composition of the Soil Bacterial Community. Front. Microbiol. 9, 1412. 10.3389/fmicb.2018.01412

63. Szukics, U., Abell, G.C.J., Hödl, V., Mitter, B., Sessitsch, A., Hackl, E., Zechmeister-Boltenstern, S., 2010. Nitrifiers and denitrifiers respond rapidly to changed moisture and increasing temperature in a pristine forest soil. FEMS Microbiol. Ecol. 72, 395–406. 10.1111/j.1574-6941.2010.00853.x

64. Takahashi, S., Tomita, J., Nishioka, K., Hisada, T., Nishijima, M., 2014. Development of a Prokaryotic Universal Primer for Simultaneous Analysis of Bacteria and Archaea Using Next-Generation Sequencing. PLOS ONE 9, e105592. 10.1371/journal.pone.0105592

65. Tang, F.H.M., Maggi, F., 2021. Pesticide mixtures in soil: a global outlook. Environ. Res. Lett. 16, 044051. 10.1088/1748-9326/abe5d6

66. Vinebrooke, R.D., Cottingham, K.L., Norberg, J., Scheffer, M., Dodson, S.I., Maberly, S.C., Sommer, U., 2004. Impacts of multiple stressors on biodiversity and ecosystem functioning: the role of species co-tolerance. Oikos 104, 451–457. 10.1111/j.0030-1299.2004.13255.x

67. Wang, F., Li, Z., Wei, Y., Su, F., Guo, H., Guo, J., Wang, Y., Zhang, Y., Hu, S., 2021. Responses of soil ammonia-oxidizing bacteria and archaea to short-term warming and nitrogen input in a semi-arid grassland on the Loess Plateau. Eur. J. Soil Biol. 102, 103267. 10.1016/j.ejsobi.2020.103267

68. White, T.J., Bruns, T., Lee, S., Taylor, J., 1990. Amplification and Direct Sequencing of Fungal Ribosomal RNA Genes for Phylogenetics. In: PCR Protocols: A Guide to Methods and Applications. Innis, M.A., Gelfand, D.H., Sninsky, J.J., White, T.J., Ed. Acad. Press., New York, NY, pp. 315–322. 10.1016/B978-0-12-372180-8.50042-1

69. Whitman, W.B., Coleman, D.C., Wiebe, W.J., 1998. Prokaryotes: The unseen majority. Proc. Natl. Acad. Sci. 95, 6578–6583. 10.1073/pnas.95.12.6578

70. Zabaloy, M.C., Garland, J.L., Gomez, M.A., 2010. Assessment of the impact of 2,4-dichlorophenoxyacetic acid (2,4-D) on indigenous herbicide-degrading bacteria and microbial community function in an agricultural soil. Appl. Soil Ecol. 46, 240–246. 10.1016/j.apsoil.2010.08.006

71. Zhang, J., Kobert, K., Flouri, T., Stamatakis, A., 2014. PEAR: a fast and accurate Illumina Paired-End reAd mergeR. Bioinformatics 30(5), 614–620. 10.1093/bioinformatics/btt593

