## Supplementary Files for "Evaluating the Effects of Environmental Disturbances and Pesticide Mixtures on N-cycle related Soil Microbial Endpoints"

### SUPPLEMENTARY TABLE AND FIGURES

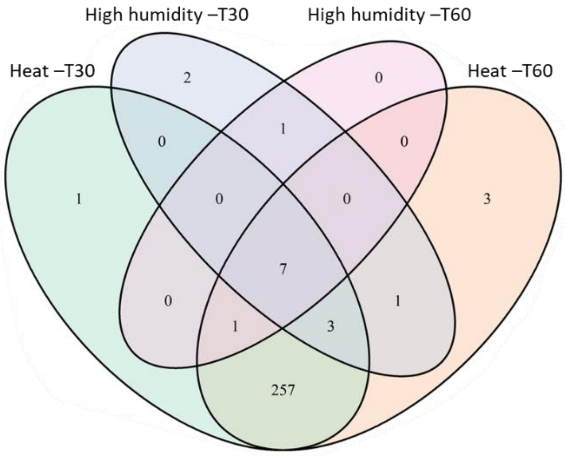


**Figure S1. Venn Diagram illustrating the number of Operational Taxonomic Units (OTUs) impacted by heat or high humidity disturbances at T30 and T60 across all pesticide treated microcosms.** OTUs whose relative abundance was significantly impacted by disturbances were identified by performing a GLM model on the most abundant 336 OTUs (abundance > 0.1% and present at least in 5/5 of the samples of one Time-A.I.-Dose-Disturbance condition). Each circle represents the set of OTUs impacted by one disturbance (high humidity or heat) at one time of sampling (T30 or T60).

**Table S1. Contribution of the microbial endpoints to the principal components of the PCA at T30 and T60.** For each axis, values represent the percentage of variance explained by each endpoint in the dataset.

| | T30 | | T60 | |
|---|---|---|---|---|
| | Axis 1 | Axis 2 | Axis 1 | Axis 2 |
| OS | 13.5 | 0.0 | 15.0 | 0.7 |
| PD | 13.3 | 0.1 | 14.5 | 1.1 |
| SR | 13.0 | 0.1 | 15.2 | 0.0 |
| 16S | 2.1 | 28.7 | 2.9 | 22.7 |
| 18S | 8.0 | 17.6 | 10.6 | 12.4 |
| ITS | 8.1 | 11.2 | 9.8 | 9.5 |
| AOA | 1.6 | 10.3 | 0.0 | 8.5 |
| AOB | 9.6 | 0.3 | 5.2 | 1.7 |
| ComaA | 7.1 | 14.7 | 6.1 | 18.8 |
| ComaB | 7.1 | 15.9 | 4.7 | 23.7 |
| $NH_4^+$ | 4.0 | 0.7 | 0.9 | 1.0 |
| $NO_3^-$ | 12.6 | 0.3 | 15.0 | 0.0 |

PD: Faith's Phylogenetic Diversity; OS: observed species; SR: Simpson's reciprocal index.
